# Differential retinal ganglion cell resilience to optic nerve injury across vertebrate species

**DOI:** 10.1101/2025.03.19.644129

**Authors:** Julie D. De Schutter, Anyi Zhang, Pieter-Jan Serneels, Lieve Moons, Luca Masin, Steven Bergmans

## Abstract

Optic neuropathies comprise a diverse group of disorders that ultimately lead to retinal ganglion cell (RGC) degeneration. Despite varying etiologies, these conditions share a conserved pathological progression: axonal damage in the optic nerve triggers progressive RGC degeneration. Understanding species-specific differences in neuronal resilience is critical for identifying key survival mechanisms and potential neuroprotective targets. In this study, we compare RGC densities and survival rates following optic nerve crush (ONC) in three vertebrate models—mice, zebrafish, and killifish—under standardized experimental conditions. Transcriptomic analysis confirmed that, similar to RBPMS in mice, Rbpms2 serves as a pan-RGC marker in zebrafish and killifish. Using these markers, we reveal significant species-specific differences in RGC density, with fish species exhibiting over a five-fold higher density than mice at equivalent life stage. Killifish also show an age-dependent decline in RGC density. Furthermore, we identify distinct injury responses across species: mice undergo rapid degeneration, losing ∼80% of their RGCs by day 14 after ONC; zebrafish maintain full RGC retention for two weeks before experiencing a loss of ∼12%; and killifish display a biphasic response to ONC, with young adults retaining two-thirds of their RGCs by day 21, while older fish exhibit a more pronounced second wave of RGC loss, ultimately preserving just over half of their RGCs by 21 days after injury. These findings highlight fundamental differences in neuroprotective capacity among species, providing a comparative framework to uncover molecular mechanisms governing RGC survival and to identify therapeutic strategies for treating optic neuropathies and neurodegeneration across diverse pathologies.

## 1. Introduction

Neuronal loss is a hallmark of both acute and chronic neurodegenerative conditions, including those affecting the visual system. Optic neuropathies encompass diverse disorders, from glaucomatous neurodegeneration to ischemic and traumatic optic nerve injuries. Despite differing etiologies, these conditions converge on a common pathological sequence: initial damage of retinal ganglion cell (RGC) axons in the optic nerve in turn triggering progressive RGC death within the retina (Ramos-Cejudo et al., 2018; Smolen et al., 2023). Being the retina’s sole output neurons, loss of RGCs disrupts visual signaling to the brain, resulting in irreversible blindness. A key challenge in treating optic neuropathies—and neurodegenerative diseases broadly—is preventing neuronal degeneration and enhancing resilience to injury. Thus, elucidating the molecular and cellular mechanisms underlying RGC vulnerability is essential for advancing neuroprotective strategies.

To this end, model organisms, and the differences among them, are essential for unraveling RGC biology and pathophysiology in retinal disorders. RGC density varies across species and age, affecting visual acuity and disease susceptibility (Baden et al., 2020). Mice are widely used due to their genetic similarity to humans and well-established models of glaucomatous injury, exhibiting a comparable central nervous system (CNS) injury response (Varadarajan et al., 2022). Zebrafish provide insights into neurorepair due to their remarkable regenerative abilities, which are maintained throughout adulthood (Van houcke et al., 2017; Marques et al., 2019). Killifish, although teleost fish, lose their regenerative capabilities at old age, recapitulating mammalian-like phenotypes after injury (Vanhunsel et al., 2022a, 2022b). Therefore, they offer a valuable model for studying age-related changes in RGC vulnerability, as shown by increased stress response systems and reduced expression of *bdnf* (brain-derived neurotropic factor) in the aged retina (Bergmans et al., 2024). Thus, species-specific differences in injury responses may uncover key factors influencing CNS neuronal survival and, in turn, identify novel neuroprotective targets that could form the foundation for new therapeutic strategies.

Although RGC survival after optic nerve crush (ONC) has been studied singularly in mice, zebrafish, and killifish, a comprehensive, cross-species analysis using whole-mount (WM) retinas for zebrafish and killifish has yet to be performed. Unlike the use of narrow field-of-view micrographs, entire retinal WMs rule out regional differences in injury response that might skew the general outcome. This is particularly relevant for killifish, which, due to their fast-growing nature, undergo significant retinal stretching during adulthood (Bergmans et al., 2023a), potentially leading to overestimation of RGC loss when using spatially pre-selected samples like cryosections (Vanhunsel et al., 2022a). Furthermore, a major limitation in fish models has been suboptimal RGC labeling methods. In zebrafish, RGC survival has largely been assessed using Tg(*isl2b*:reporter) lines (Skarie et al., 2015; Skarie and Nickells, 2016; Chen et al., 2022). Recent single cell RNAseq data however disputes the validity of *isl2b* (ISL LIM homeobox 2b) as a pan-RGC marker in zebrafish, as its expression is neither uniform nor ubiquitous across RGC subtypes (Kölsch et al., 2021). In killifish, RGCs have been labelled via retrograde biocytin tracing (Vanhunsel et al., 2022a; Bergmans et al., 2023a), but this can lead to incomplete labelling of RGCs when the procedure is not carried out correctly. Furthermore, RGC axons are also marked by retrograde biocytin tracing, occluding the view of the ganglion cell layer in regions of high axonal density, such as in the central retina. Conversely, Rbpms2 (RNA-binding protein with multiple splicing 2) has emerged as a pan-RGC marker in both species based on RNA sequencing data (Kölsch et al., 2021; Bergmans et al., 2024), but has yet to be used to assess RGC densities and/or survival rates following ONC in these teleost models.

In this study, we perform optic nerve crush (ONC) in three key model organisms—mice, zebrafish, and killifish—to directly compare retinal ganglion cell (RGC) survival across species. By standardizing experimental conditions within a single laboratory, we minimize inter-laboratory variability, strengthening cross-species comparisons. Using pan-RGC markers, we reveal species-specific RGC densities and injury responses. The studied fish species exhibit significantly higher RGC densities compared to mice of an equivalent life stage, and RGC density declines with increasing age in killifish. Furthermore, murine RGCs exhibit low resilience to ONC, zebrafish preserve RGC numbers for two weeks before a delayed degenerative phase, and killifish display a biphasic cell loss pattern affected by age. As such, our findings provide a foundation for future studies exploring molecular mechanisms underlying differential RGC resilience across vertebrates.

## 2. Material and Methods

### 2.1 Transcriptomic data

To display the expression pattern of RGC markers across the retina, previously published scRNAseq datasets were used: 1) Li et al. for mouse (Li et al., 2024), 2) Hoang et al. for zebrafish (Hoang et al., 2020), and 3) Bergmans et al. for killifish (Bergmans et al., 2024). All three datasets were used “as is” and the original dimensionality reduction projections and clustering were used for plotting. For zebrafish, only the non-injured cells of the NMDA dataset of Hoang et al. were used.

### 2.2 Animal housing

Mice (*Mus musculus*) of the C57Bl/6N strain (Charles River Laboratories, France) were housed under standard laboratory conditions (12-hour light/12-hour dark cycle, 21°C, 50% humidity) (Masin et al., 2021). Water was accessible at all times and they were fed ad libitum. All experiments were carried out using 10-week-old mice of both sexes.

Zebrafish (*Danio rerio*) of the AB wildtype strain were raised and maintained under standard laboratory conditions as described (Beckers et al., 2021), specifically at 28°C with a conductivity of 650 µS and pH of 7.5 on a 14-hour light/10-hour dark cycle. They were fed twice a day with a mixture of dry food and brine shrimp (Artemia *Salina nauplii*, Ocean Nutrition). All experiments were conducted on 21-week-old adult zebrafish of similar size, including both males and females.

African turquoise killifish (*Nothobranchius furzeri*) were raised and housed at standard laboratory conditions (Bergmans et al., 2023a), i.e., at 28°C with a conductivity of 600 µS and pH of 7.0 on a 12-hour light/dark cycle. Fish were fed twice a day with a combination of brine shrimp and *Chironomidae* larvae. All experiments made use of 6- and 18-week-old killifish of the GRZ-AD inbred strain. Only female fish were used to minimize sex difference as male and female killifish differ substantially in size and age trajectory, potentially altering both cell densities and survival properties.

All animal experiments received approval from the KU Leuven Animal Ethics Committee and were conducted in strict compliance with the European Communities Council Directive of 20 October 2010 (2010/63/EU).

### 2.3 Optic nerve crush

In mice, a unilateral ONC was performed as previously described (De Groef et al., 2016; Masin et al., 2021). Briefly, animals were anesthetized through intraperitoneal injection of ketamine (75 mg/kg body weight, Nimatek, Eurovet) and medetomidine (1 mg/kg, Domitor, Pfizer). Next, the optic nerve was exposed after an incision of the conjunctiva and crushed 1 mm from the optic nerve head using a Dumont #7 cross-action forceps (Fine Science Tools) for 10 seconds. After ONC, anesthesia was reversed using atipamezole (1 mg/kg, Antisedan, Pfizer). Additionally, local analgesia (oxybuprocaïne 0.4%, Unicaïne) was applied to the eye before surgery, and antibiotic ointment (tobramycin 0.3%, Tobrex) was applied afterward.

Detailed protocols to unilaterally crush both zebrafish and killifish optic nerves have been previously reported (Beckers et al., 2023; Vanhunsel et al., 2023). Briefly, fish were anesthetized using 0.03% Tris-buffered tricaine (MS-222, Sigma-Aldrich). The connective tissue surrounding the eye was removed and the eyeball was lifted out of its orbit, exposing both the optic nerve and ophthalmic artery. Using a Dumont #7 cross-action forceps (Fine Science Tools), the optic nerve was crushed at a distance of 0.5 mm from the optic nerve head for 10 seconds, without damaging the ophthalmic artery. Both the distance from the optic nerve head and the duration of the crush were kept constant for young adult zebrafish, young adult and old killifish. After the procedure, the fish were returned to system water to recover.

### 2.4 Retrograde labelling of fish retinal ganglion cells using biocytin

Retrograde labelling of both teleost RGCs was performed as described in detail for both zebrafish (Beckers et al., 2023), and killifish (Vanhunsel et al., 2023). Fish were anesthetized and the eye was lifted from its socket as described above. The optic nerve was completely cut at a distance of 500 µm from the optic nerve head. A gel foam, drained with a saturated biocytin solution, was placed at the proximal end of the optic nerve. The fish were awakened in a recovery tank for 3 hours to allow passive retrograde transport of the tracer. Next, fish were euthanized, and tissues were collected (see 2.5 Tissue sampling and processing).

### 2.5 Tissue sampling and processing

All mice were killed with an overdose of sodium pentobarbital (60 mg/kg, Dolethal, Vetoquinol), while fish were euthanized using an overdose of tricaine (0.1% Tris-buffered tricaine, MS-222). Animals were subsequently transcardially perfused using 0.9% NaCl (mice) or phosphate buffered saline (PBS, fish) and 4% paraformaldehyde (PFA) as previously described (Wu et al., 2021; Mariën et al., 2022; Bergmans et al., 2023b). Eyes were fixated in 4% PFA for 1 hour after enucleation and subsequently washed 3 times with PBS and stored in storage buffer (0.4M NaN_3_ in PBS) until further use. Retinas were dissected and whole-mounted, fixated for an additional hour in 4% PFA, washed 3 times in PBS and stored in storage buffer until further use.

Next, retinal WMs were immunohistochemically stained for RBPMS (mice) or Rbpms2 (fish). WMs were permeabilized by washing steps in 0.5% Triton X-100 in PBS (PBST 0.5%) and a 15 min freeze-thaw step at –80°C. Samples were blocked for 2 hours using pre-immune donkey serum (PID, 1:5 in PBST 2%). Hereafter, retinal WMs were incubated with the primary antibody overnight in PBS with 2% Triton X-100 and 10% PID. Details on primary antibodies are listed in Table 1. Tissues were washed several times with PBST 0.5% before an incubation of 2 hours with the secondary antibody (Alexa-647-conjugated donkey anti-rabbit IgG, 1:200, Thermofisher). Retinas from fish that were retrogradely traced using biocytin were incubated with streptavidin-Alexa488 for 2 hours. All steps were carried out at room temperature. Retinas were mounted on glass slides with anti-fading mounting reagent Mowiol (10%, Sigma-Aldrich).

**Table 1:** List with primary antibodies used for immunohistochemistry.

| Target species | Antibody | Manufacturer | Catalog number | Host species | Dilution |
| --- | --- | --- | --- | --- | --- |
| Mice | Anti-RBPMS | PhosphoSolutions | AB_2492225 | Rabbit | 1:250 |
| Zebrafish | Anti-Rbpms2 | Abcam | Ab181098 | Rabbit | 1:200 |
| Killifish |  |  |  |  |  |

Finally, mosaic images of entire retinal WMs were imaged using a wide-field epifluorescent microscope (Leica DM6) containing a HC PL FLUOTAR L 20x/0.40 CORR objective (resolution = 2.17 pixels/µm).

### 2.6 Retinal whole-mount analysis

#### 2.6.1 Establishment of automated counting models

The deep learning-based cell counting models for zebrafish and killifish RGCs were derived from RGCode (Masin et al., 2021) via transfer-learning. Briefly, frames of fixed size (177 by 177 µm) were obtained from Rbpms2-stained retinal WMs, sampling the central, mid and peripheral retina equally. The dataset was divided into training and testing datasets, and the cells were annotated by two independent expert counters. The size and number of frames per dataset is reported in Table 2. Both counters annotated all testing frames, while the training ones were equally split between the two counters. Transfer-learning was performed starting from the weights of the original RGCode neural network, as described previously (Masin et al., 2021). The performance of the newly-trained models was evaluated by linear-regression analysis against the human-annotated frames of the testing datasets. If the slope and coefficient of determination of the model were comparable to those between human counters, the performance was deemed satisfactory. Performance metrics of the final models are reported in Figure S1.

**Table 2:** Composition of training datasets for zebrafish and killifish.

| Species | Dataset | N frames |
| --- | --- | --- |
| Zebrafish | Train | 48 |
|  | Test | 36 |
| Killifish | Train | 36 |
|  | Test | 24 |

#### 2.6.2 Analysis

The number and density of RGCs per retina, together with the retinal area, were obtained by running the RGCode2 pipeline. Murine retinas were analyzed using the original RGCode model, while for zebrafish and killifish the newly-trained models were used. For retinal segmentation, the original RGCode model was used for all organisms, as its performance was satisfactory across all three animal species. RGCs density across the whole retina was obtained by dividing the total number of RGCs by the retinal area. Isodensity maps of the retina were generated as probability density functions via gaussian kernel density estimation (KDE). The KDE was generated from the centroids of the detected cells, with a bandwidth of 100 µm, and scaled to represent cells per square millimeter.

### 2.7 Statistical analysis

All data analyses were performed on raw, unsaturated micrographs. For visualization, some images were contrast-enhanced by adjusting the white point, applying identical, linear enhancements across comparable images. Statistical analyses included ANOVA, t-tests and U-tests, as described in the figure legends. The median with Kruskal– Wallis ANOVA and Mann-Whitney U-tests were used when data failed the Shapiro–Wilk normality test; otherwise, the mean with Welch ANOVA was chosen. All data processing, plotting, statistical analyses, were performed in Python using pandas, seaborn, matplotlib, and dabest (Ho et al., 2019). A p-value < 0.05 was considered significant.

## 3. Results

### 3.1 Validation of Rbpms2 as a pan-RGC marker in the teleost retina

RBPMS is a widely recognized pan-RGC marker in the mouse retina and it has replaced BRN3A (*Pou4f1*, POU class 4 homeobox 1) several years ago (Figure 1A-C) (Rodriguez et al., 2014; Rheaume et al., 2018; Tran et al., 2019; Masin et al., 2021; Li et al., 2024). In contrast, *rbpms2b* has only recently been reported as a potential pan-RGC marker in zebrafish and killifish based on single cell RNA sequencing (Figure 1D,E,G,H) (Kölsch et al., 2021; Bergmans et al., 2024). While *isl2b* has been traditionally used for labelling zebrafish RGCs, it is not ubiquitously expressed in RGCs of zebrafish or killifish (Figure 1F,I) (Hoang et al., 2020; Kölsch et al., 2021; Bergmans et al., 2024). To validate Rbpms2 as a pan-RGC marker for teleosts, we performed retrograde labelling assays known to label all RGCs in zebrafish and killifish. We observed near-complete co-labeling of biocytin and Rbpms2 in the zebrafish (Figure 1J) and killifish (Figure 1K) retina. However, biocytin labeling was heterogenous, with intensities ranging from faint to bright, whereas Rbpms2 provided uniform labeling of all RGCs. Additionally, Rbpms2 specifically marked RGC somas, unlike biocytin, which also labeled axons, masking the retinal ganglion cell layer and complicating quantification.

**Figure 1:**
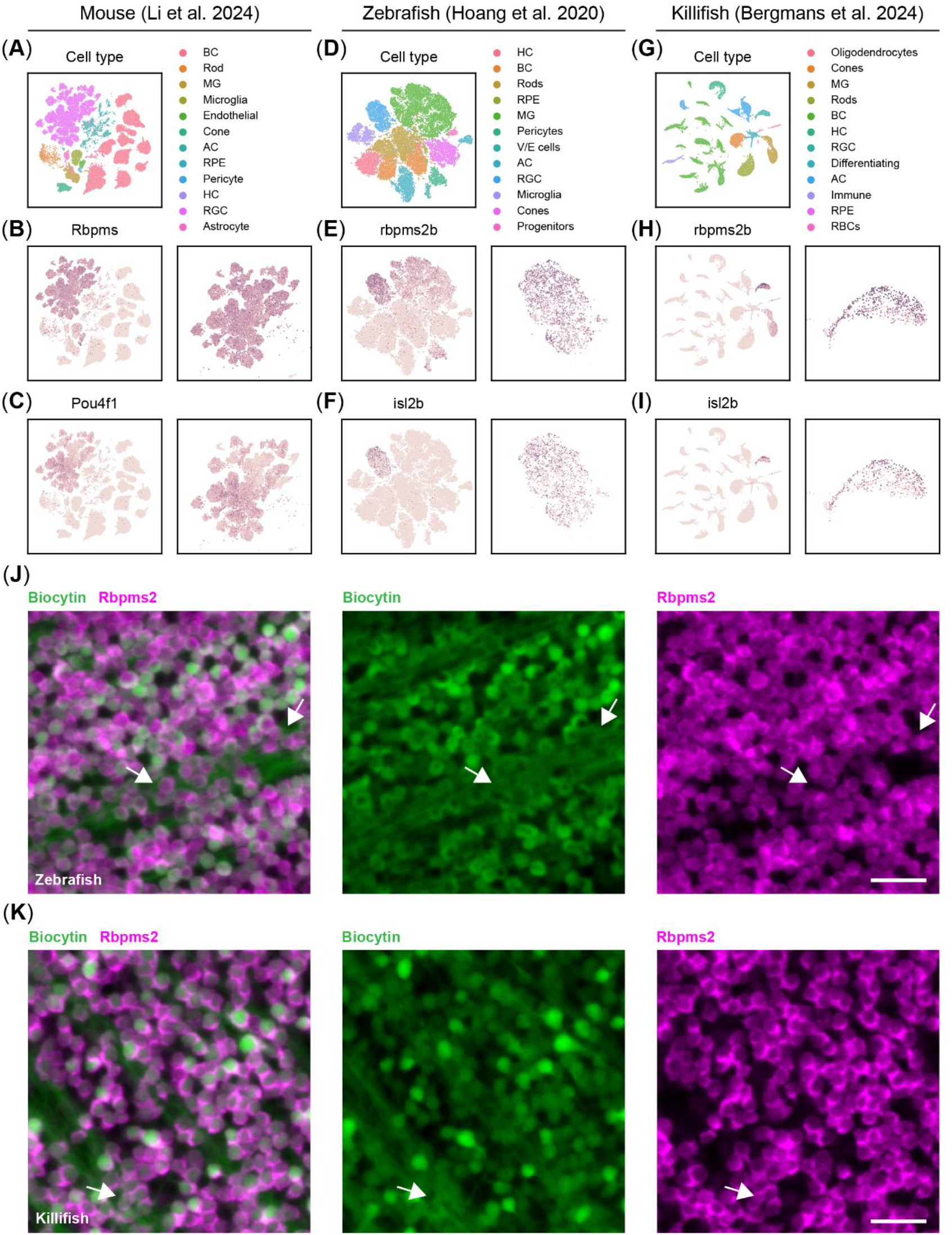
Validation of Rbpms2 as a pan-RGC marker in the teleost. **(A)** UMAP projection of the Li et al. 2024 dataset, showing the clusters of major retinal cell types in the adult mouse retina. **(B)** Rbpms is a pan-RGC marker showing homogenous expression across all RGCs in the murine retina (zoom on the RGC cluster on the right). **(C)** Contrary to Rbpms, Pou4f1, also known as Brn3a, does not show homogenous expression across all RGCs in mice. **(D)** tSNE projection of the Hoang et al. 2020 dataset (displaying only non-injured cells) revealing the major retinal cell types in the adult zebrafish retina. **(E)** rbpms2b is a pan-RGC marker showing homogenous expression across all RGCs in the adult zebrafish retina (zoom on the RGC cluster on the right). **(F)** isl2b, the most commonly used promoter in transgenic reporter lines for RGCs in zebrafish, does not show homogenous expression across all RGCs in adult zebrafish. **(G)** UMAP projection of the Bergmans et al. 2024 dataset, showing the clusters of major retinal cell types in the adult killifish retina. **(H)** As in zebrafish, rbpms2b emerges as the most homogenous marker for RGCs in the adult killifish retina (zoom on the RGC cluster on the right). **(I)** isl2b, like in zebrafish, does not show homogenous expression across all RGCs in adult killifish. **(J)** Representative micrographs of zebrafish retinal WMs, in which the RGCs are retrogradely traced with biocytin and immunostained for Rbpms2. Contrary to biocytin, Rbpms2 staining results in more homogenous, somatic labelling and labeled cells are not occluded by axonal bundles in the nerve fiber layer (arrows). Scale bar 25 µm. **(K)** Representative micrographs of killifish retinal WMs, in which the RGCs are retrogradely traced with biocytin and immunostained for Rbpms2. As in zebrafish, Rbpms2 staining results in homogeneous somatic labeling, while biocytin tracing is more heterogenous and RGCs are occasionally occluded by axon bundles (arrows). Scale bar 25 µm. Abbreviations: AC, amacrine cell; BC, bipolar cell; HC, horizontal cell; MG, Müller glia; RBC, red blood cell; RGC, retinal ganglion cell; RPE, retinal pigment epithelium; V/E, vascular/endothelial; WMs, whole-mounts

Thus, both RBPMS and Rbpms2 serve as reliable markers for assessing RGC densities and survival in murine and teleost retinas, respectively, and can be leveraged for automated cell counting platforms such as RGCode (Masin et al., 2021).

### 3.2 Comparative analysis of RGC density in the retina of mice, zebrafish and killifish

RGC densities were compared across species, including young adult mice (C57Bl/6N, 10-week-old), young adult zebrafish (AB, 21-week-old), and African turquoise killifish (GRZ-AD), a teleost gerontology model. In killifish, both young adult (6-week-old) and old (18-week-old) age groups were analyzed to assess age-related changes in RGC density. To this end, we developed RGCode2, an expansion of RGCode (Masin et al., 2021), a RGC counting platform that, next to murine RBPMS- and FluoroGold-positive RGCs, has been trained to count Rbpms2-positive RGCs in retinal WMs of both zebrafish and killifish. Furthermore, RGCode2 is able to segment both murine and teleost retinas, allowing to determine retinal areas and infer RGC densities.

Scaled images of retinal WMs from mice, zebrafish and killifish reveal significant differences in retinal size (Figure 2A-D), confirmed by retinal area analysis based on automated segmentation by RGCode2 (Figure 2I). Notably, killifish exhibit substantial retinal expansion between 6 and 18 weeks (Figure 2C,D,I), consistent with pervious findings (Bergmans et al., 2023a). Additionally, a clear difference in individual RGC soma size between mice and the studied teleost species is visually evident in Figure 2E-H. On average, 10-week-old mouse retinas contain 44,499 ± 470 RGCs (Figure 2J), as previously reported (Masin et al., 2021). This is significantly fewer than zebrafish and killifish at comparable life stages (21- and 6-week-old, respectively), which have on average 66,262 ± 2419 and 74,597 ± 2,122 RGCs in their total retina (Figure 2J). Due to the lifelong growth of fish, including zebrafish and killifish (Van houcke et al., 2019; Vanhunsel et al., 2021; Bergmans et al., 2023a), RGC numbers increase significantly with age in killifish, reaching an average of 132,159 ± 3,477 RGCs in 18-week-old killifish (Figure 2J).

**Figure 2:**
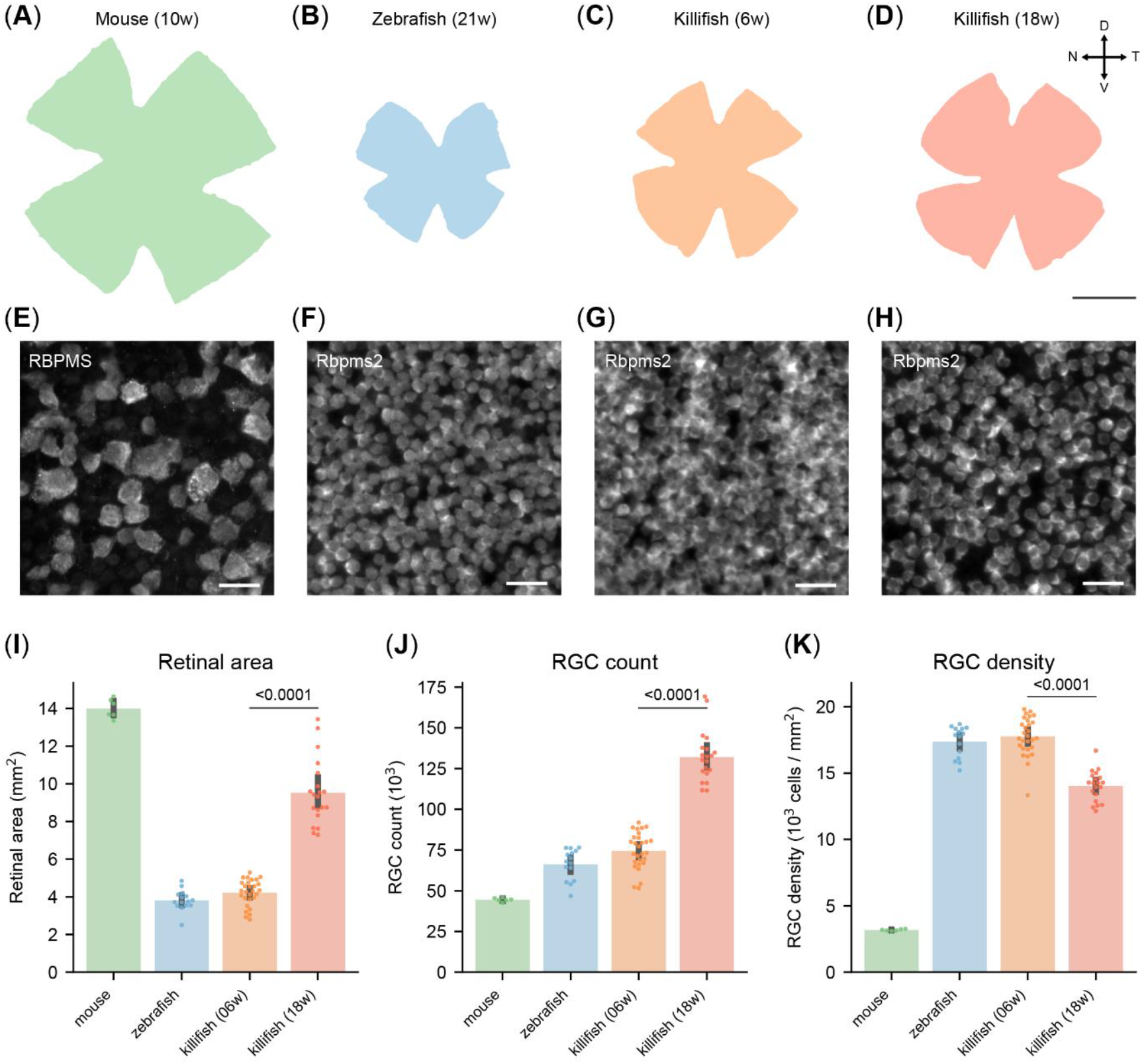
Different RGC density in retinas of adult mice, zebrafish and killifish. (**A-D**) Scaled representation of retinas from young adult mice (10-week-old, **A**), young adult zebrafish (21-week-old, **B**), young adult killifish (6-week-old, **C**) and old killifish (18-week-old, **D**). Mice have larger retinas than zebrafish and young killifish, while the retina of aged killifish is considerably larger than that of their younger counterparts. Scale bar 1mm. (**E-H**) Representative micrographs of RGCs labelled with RBPMS (mouse, **E**) or Rbpms2 (fish), sampled from the temporal retina. Young adult zebrafish (**F**) and killifish (**G**) show a comparable density, higher than the one of old killifish **(H)** and young adult mice. Moreover, RGCs from the fish species are considerably smaller than the ones of mice. Scale bar 25 µm. **(I)** Automated quantification of the area of retinal WMs, revealing that unlike young adult fish, old killifish approach the size of murine retinas. **(J)** Automated quantification of RGC numbers in retinal WMs. Mice exhibit the lowest RGC count, with approximately 45,000 cells. In contrast, young fish possess around 70,000 RGCs. Aged killifish have the highest count, reaching approximately 125,000, nearly twice as many as young killifish. **(K)** Automated quantification of RGC density in retinal WMs. RGC density is considerably lower in mice compared to fish species. Notably, old killifish exhibit a significantly reduced RGC density compared to young adult fish. Abbreviations: D, dorsal; N, nasal; RGCs, retinal ganglion cells; T, temporal; V, ventral; WMs, whole-mount

RGC densities were calculated to reduce the impact of dissection artifacts and function as a more reliable interspecies comparison metric. RGC densities also serve as a proxy for visual acuity since it determines the degree of spatial detail that can be transmitted to the brain. Young adult zebrafish (21-week-old) and killifish (6-week-old) exhibit similar RGC densities of 17,390 ± 301 and 17,769 ± 272 RGCs/mm^2^, respectively, which are over five times higher than in young adult mice (3,179 ± 26 RGCs/mm^2^, Figure 2E,F,G,K). In old killifish (18-week-old), density declines to 14,041 ± 255 RGCs/mm^2^ (Figure 2H,K), but remains more than four times higher than in 10-week-old mice. This reduction in RGC density in old killifish is attributed to age-related tissue stretching, where the distance between cell centroids increases over time (Bergmans et al., 2023a). Scaled isodensity projections (Figure S2) confirm previously published density maps of mice (Masin et al., 2021) and zebrafish (Mangrum et al., 2002). Killifish exhibit an RGC distribution similar to zebrafish, with the highest density in the ventrotemporal quadrant of the retina (Figure S2). Average retinal area, RGC counts and densities of the three studied species are reported in supplementary Table S1.

Although the retina is highly conserved across vertebrates in terms of anatomy, including its layered structure and cell types, RGC densities vary greatly between species. These differences highlight the importance of cross-species comparisons of RGC function and responses to stimuli or injury, providing valuable insights into neural network dynamics and injury mechanisms.

### 3.3 Differential RGC susceptibility to optic nerve injury in canonical vertebrate models

Extensive literature describes a striking difference in intrinsic survival capacity of RGCs between mammals and teleost fish (Zou et al., 2013; Lukowski et al., 2019; Tran et al., 2019; Masin et al., 2021; Vanhunsel et al., 2022a). To validate these findings under standardized conditions, we conducted a comparative study within a single laboratory, using a consistent ONC injury model across the investigated species. This approach eliminates interlaboratory technical variation, allowing for a direct comparison of RGC survival and resilience in mice and zebrafish, two canonical vertebrate models.

Based on a previously published RGC survival curve in mice (Tran et al., 2019), we selected two key timepoints to assess RGC survival upon ONC across species (Figure 3A). At 7 days post injury (dpi) RGC loss is actively ongoing, while by 14dpi RGC death reaches a plateau with nearly all susceptible RGCs lost (Tran et al., 2019; Masin et al., 2021). In our study, 59% [IQR:58%, 62%] of the RGCs had died in mice by 7dpi, consistent with previous reports (Masin et al., 2021), and by 14dpi, only 19% [IQR: 18%, 19%] remained (Figure 3B,D). This severe degeneration underscores the mammalian CNS’s limited ability to withstand injury. In contrast, adult zebrafish, renowned for their astonishing regenerative abilities (Marques et al., 2019), display strong injury resilience, with 97% [IQR: 95%, 100%] of their RGCs surviving the first 14 days after ONC (Figure 3C,E). However, recent observations after optic nerve transections indicate late RGC degeneration in the adult zebrafish retina (Zou et al., 2013). As such, we also investigated a later timepoint after ONC (Figure 3A) to assess whether this late loss of RGCs also occurs after ONC. Similarly to optic nerve transection, zebrafish do present with a minor, but significant, decrease in their RGC survival 21dpi, with 88% [IQR: 87%, 91%] of the RGCs surviving (Figure 3C,E). This minor loss might, however, be negligible for visual function as zebrafish are known to regain primary visual abilities between 10 and 15dpi (Van houcke et al., 2017), and further refine neuronal circuits to fully recover complex visual behaviors between 30 and 50dpi (Kaneda et al., 2008). Of note, although zebrafish are ever growing organisms, their retinas did not exhibit any significant expansion in area during the evaluated time window (Figure S3A), nor do they present with alterations in RGC density during adulthood (Van houcke et al., 2019).

**Figure 3:**
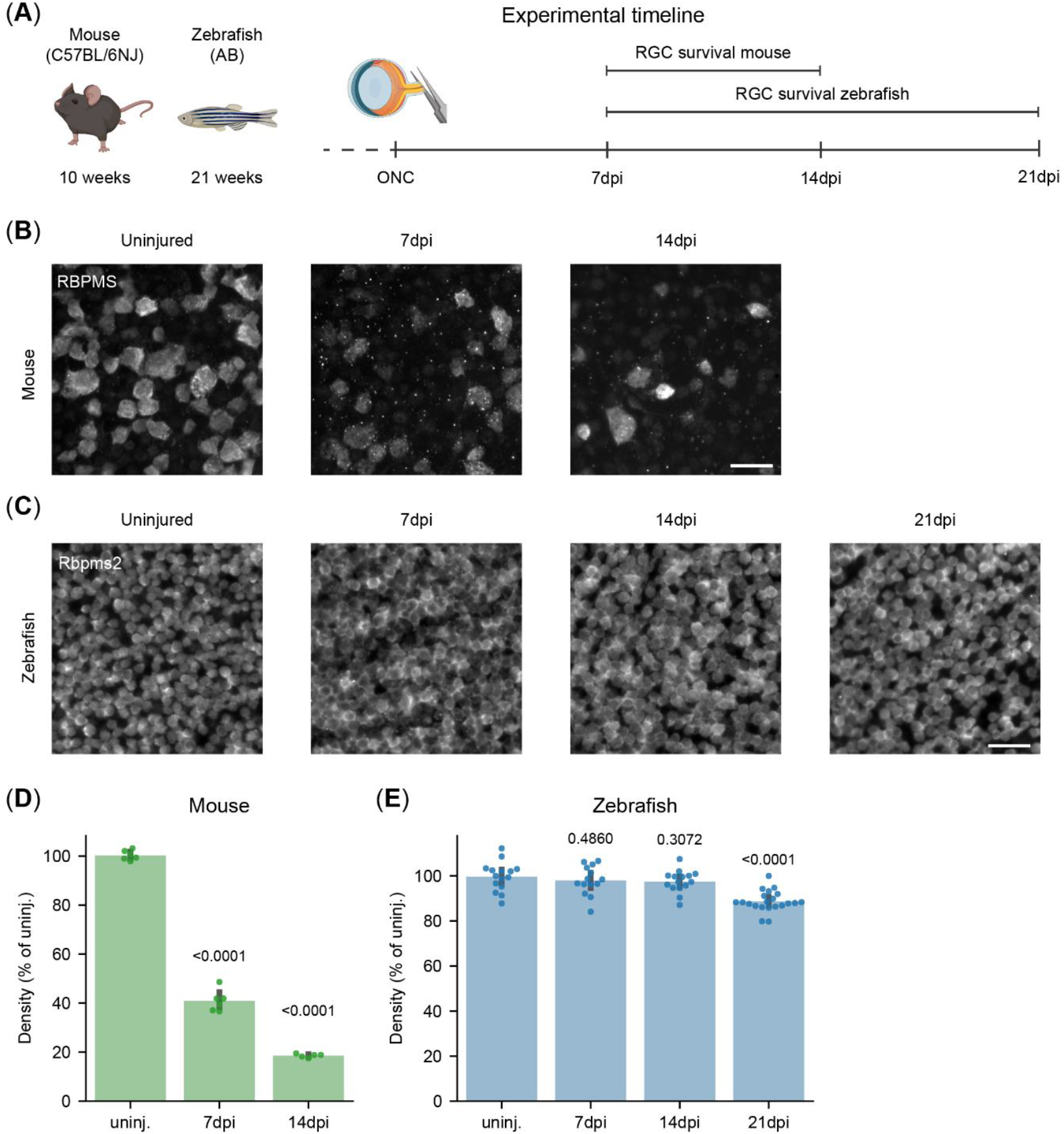
Differential RGC susceptibility to optic nerve injury in canonical vertebrate models. **(A)** Experimental timeline for the RGC survival experiment in young adult mice and zebrafish. RGC survival was evaluated at 7 and 14 days post ONC in mice and at 7, 14 and 21dpi in zebrafish. With assets from BioRender.com. **(B)** Representative micrographs of RPBMS-stained murine retinal WMs following ONC injury show substantial RGC loss at 7 dpi, which becomes even more pronounced by 14 dpi. Scale bar 25 µm. **(C)** Representative micrographs of Rpbms2-stained zebrafish retinal WMs following ONC injury. ONC injury leads to no appreciable loss of RGCs at 7 and 14dpi, but a minor loss can be observed at 21dpi. Scale bar 25 µm. **(D)** Quantification of RGC survival in adult murine WMs. ONC leads to the loss of over 50% of RGCs at 7dpi, and a further one until 14dpi, when only about 20% of the RGCs remain. **(E)** Quantification of RGC survival in adult zebrafish WMs. There is no significant loss of RGCs within the first two weeks after ONC. A small but significant decrease is measured at 21dpi, with approximately 10% of the RGCs lost. Data from 2 independent experiments, presented as percentages relative to the median of uninjured retinas and presented as median ± 25-75^th^ CI. Kruskal-Wallis ANOVA. P-values reported within the figure. Abbreviations: CI, confidence interval; dpi, days post injury; ONC, optic nerve crush; RGCs, retinal ganglion cells; WMs, whole-mounts

To conclude, we confirm the distinct resilience profiles of these two canonical vertebrate models, with mice displaying minimal neuroprotection and zebrafish exhibiting a significantly higher neuroprotective capacity.

### 3.4 Biphasic RGC loss in killifish after optic nerve crush injury

To assess the impact of aging on RGC survival, we used the African turquoise killifish, a well-established model in gerontology. Retinal aging in this species mirrors key hallmarks of human aging, including increased oxidative stress, stem cell exhaustion, gliosis, and inflammaging (Vanhunsel et al., 2021; Bergmans et al., 2024). Furthermore, neuroregenerative capacity declines with age, as older killifish fail to recover from CNS injury, unlike their younger counterparts (Van houcke et al., 2021; Vanhunsel et al., 2022a). Accordingly, we examined RGC survival in young adult (6-week-old) and old (18-week-old) killifish (Figure 4A).

**Figure 4:**
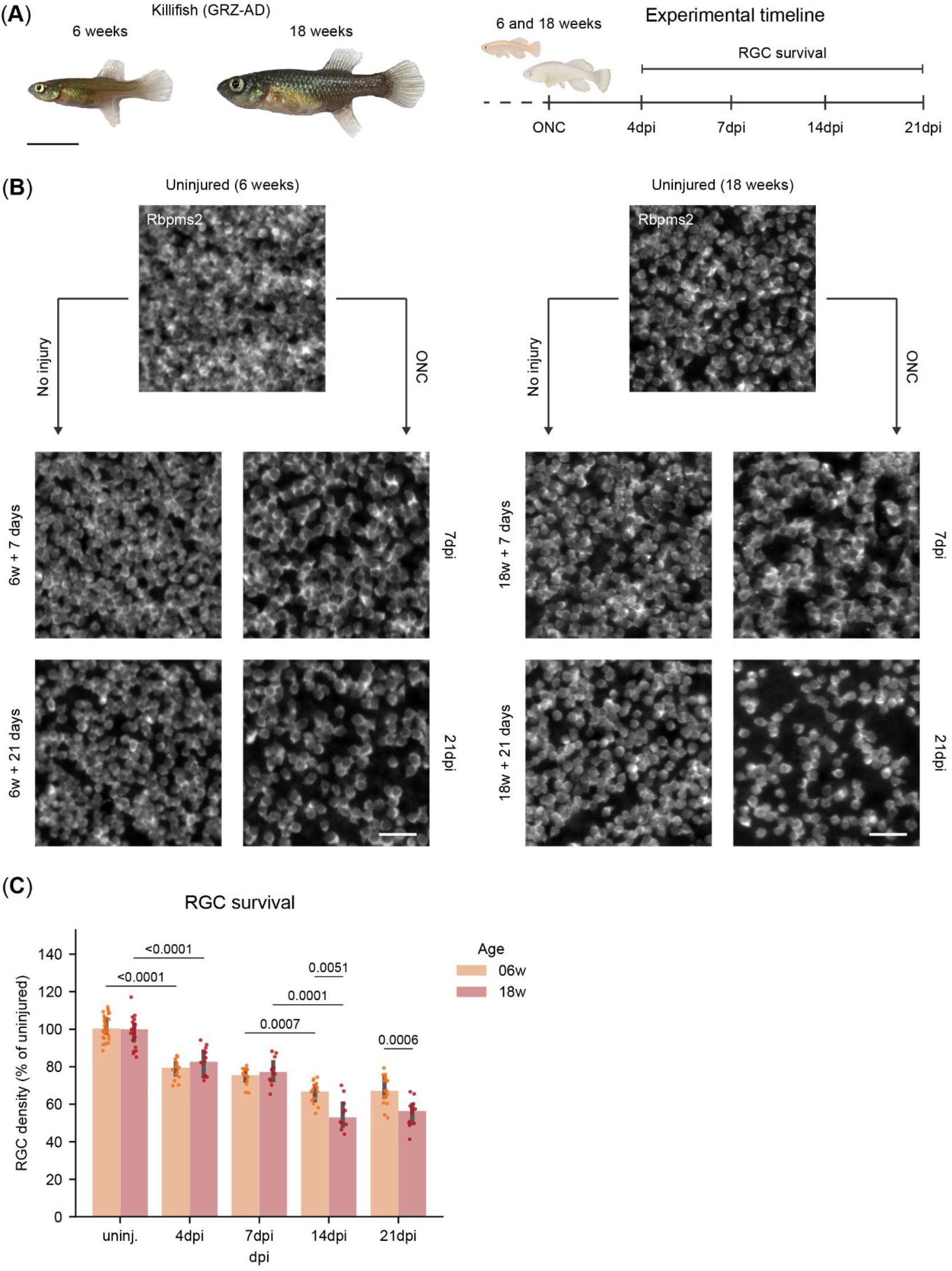
Biphasic RGC loss in killifish after optic nerve crush injury. **(A)** Representative image of a young adult (6-week-old) and old killifish (18-week-old) and experimental timeline for the RGC survival experiment, where RGC survival is evaluated at 4, 7, 14, and 21 days following ONC injury. Scale bar: 1 mm. **(B)** Representative micrographs of Rbpms2-stained WMs of young adult (6-week-old) and old killifish (18-week-old). For both ages, an appreciable loss of RGCs is evident at 7dpi with further loss at 21dpi, when compared to uninjured age-matched control fish. **(C)** Quantification of RGC survival in adult killifish WMs shows a first wave of RGC loss at 4 dpi, with 20% of the RGCs lost in both age groups, and this loss remains steady through 7 dpi. A second wave of loss is observed at 14 dpi, with older fish losing more RGCs (50%) compared to young fish (40%). No further loss is detected at 21 dpi in either age group. Data from 2 independent experiments, presented as percentages relative to the median of their uninjured age-matched control and presented as median ± 25-75th CI. Two-way Kruskal-Wallis ANOVA with pairwise Mann-Whitney U tests. P-values reported within the figure. Abbreviations: CI, confidence interval; dpi, days post injury; ONC, optic nerve crush; RGCs, retinal ganglion cells; WMs, whole-mounts

The killifish retina, however, undergoes rapid expansion, particularly during early life stages, due to its explosive growth (Vanhunsel et al., 2021; Bergmans et al., 2023a). This growth is driven by a dynamic balance between cell addition and tissue stretching, a process extensively studied by Bergmans and colleagues (Bergmans et al., 2023a). To account for this rapid growth in our experimental time window, we focused on adjusting for retinal expansion when assessing RGC density and survival. As expected, young adult killifish show significant retinal expansion between 6 weeks and 6 weeks plus 21 days (Figure S3B), while this was not the case in old killifish (18 weeks until 18 weeks plus 21 days) (Figure S3C). Next, when evaluating RGC densities in young adult killifish, we indeed also observe a slight increase from 6 weeks to 6 weeks plus 7 days, followed by a decline in RGC density over the next two weeks, which becomes significant at 6 weeks plus 21 days (Figure S3D). This alteration in RGC density profile over time is not observed in old killifish (Figure S3E). To account for the significant differences in cellular densities in young adult killifish, due to retinal growth across the lifespan of the killifish (Figure S3F-H), we opted to normalize RGC densities after injury to uninjured age-matched control fish (AMCs).

In contrast to zebrafish, both young adult and old killifish exhibit a rapid loss of RGCs following ONC, with approximately 79% [IQR: 75%, 82%] and 83% [IQR: 75%, 88%] of RGC surviving at 4dpi, respectively, a level that remains stable until 7dpi (Figure 4B,C). This initial phase of RGC loss mirrors the pattern observed in mice (Tran et al., 2019). After the first week, a second wave of cell death occurs, reducing RGC survival in young adult killifish to 67% [IQR: 63%, 71%] by 14dpi, after which it remains stable through 21dpi (Figure 4B,C). Intriguingly, while the magnitude of cell death is similar between 6- and 18-week-old killifish during the first wave, it differs during the second wave. Here, RGCs in old killifish display a higher vulnerability compared to those in younger animals, resulting in a loss up to 44% [IQR: 41%, 50%] (Figure 4B,C). Average retinal area, RGC counts and densities of the three studied species after injury are reported in supplementary Table S2 while statistical details comparing young-adult and old killifish are reported in supplementary Table S3.

These findings indicate that while both age groups exhibit a biphasic pattern of RGC loss, aging is a key factor determining the overall extent of degeneration, primarily due to the increased RGC loss observed during the second wave.

## 4. Discussion and conclusion

In this study, we validated the use of Rbpms2 as a pan-RGC marker to study ganglion cell numbers in both zebrafish and killifish. Additionally, we present an updated version of the automated RGC counting platform, RGCode (Masin et al., 2021), trained to count Rbpms2-positive RGCs of teleost species. Using RGCode2, we demonstrate that mice exhibit significantly lower RGC densities than zebrafish and killifish at equivalent life stages and that in killifish RGC density declines with age (Bergmans et al., 2023a). We further evaluated RGC survival following ONC across the three species. Consistent with previous reports, mice retained only one-fifth of their RGCs by 14dpi (Tran et al., 2019; Masin et al., 2021; Claes and Moons, 2022). Zebrafish exhibited no significant RGC loss within the first two weeks post injury, as previously reported (Zou et al., 2013), but presented with a moderate yet significant decline of approximately 10% by 21dpi. Killifish displayed a biphasic response; at 21dpi, young adults retained two-thirds of their RGCs, whereas older individuals experienced a more pronounced decline, preserving just over half their RGCs.

Despite the conserved cytoarchitecture of the vertebrate retina, from lampreys to humans (Ramon y Cajal, 1893), interspecies differences are evident. These variations include number of neuronal (sub)types, such as photoreceptor diversity, as well as species-specific transcriptional profiles within neuronal classes (Hahn et al., 2023). In a comprehensive review, Baden (2024) explored vertebrate retinal evolution, demonstrating that while retinal circuits are fundamentally conserved, species-specific adaptations arise based on ecological niche, visual demands and evolution (Baden, 2024b, 2024a). This is often linked to the diversity of photoreceptors that have evolved within an ecological niche, for example to be suited for nocturnality (less types) or diurnally (more types). Clades with higher photoreceptor types, such as fish and birds, often present circuitry specialized for specific behaviors, e.g. UV vision for prey-capture in fish. Therefore, to accommodate these circuits, Baden argues that a higher diversity of photoreceptors is linearly correlated to a higher density of RGCs in the inner retina (Baden, 2024b, 2024a). Our study further supports Baden’s hypothesis, as we observed significantly higher RGC densities in killifish and zebrafish, which possess five photoreceptor types, as compared to mice, which only possess three.

RGC survival in the mouse retina has been extensively studied, demonstrating a sigmoidal monophasic loss pattern during the first three months following injury (Tran et al., 2019; Yang et al., 2020). Initially, RGC loss is minimal within the first 3 days, followed by a rapid decline of approximately 70% over the next 5–7 days, ultimately stabilizing at around 10% survival by 30dpi (Tran et al., 2019; Yang et al., 2020). Tran and colleagues identified 45 RGC subtypes in mice and classified them based on injury susceptibility into resilient (8.1% of the total population), intermediate (27.2%), and susceptible (64.7%) groups. Resilient RGCs decline gradually, with 50% survival at 14dpi, whereas intermediate and susceptible populations undergo peak cell death at 4– 7dpi and 3–4dpi, respectively (Tran et al., 2019). A similar sigmoidal monophasic loss pattern has been observed in rats following optic nerve injury (Levkovitch-Verbin et al., 2003; Guo et al., 2020). Studies using partial optic nerve transection in rats further suggest that this characteristic monophasic loss profile results from co-occurring primary and secondary injuries responses (Guo et al., 2020). Whereas primary degeneration is typically defined as the injury resulting directly from the initial lesions, secondary degeneration is described by neuronal loss as a consequence to the primary injury (Oyinbo, 2011). As such, secondary degeneration is typically a consequence of accumulation of toxic factors such as reactive oxygen species, calcium release into the extracellular space by dying cells, glutamate excitotoxicity, and (peripheral) immune cell (over)activation, eventually resulting in cell death (Park et al., 2004; O’Hare Doig et al., 2014; Stirling et al., 2014; Jones et al., 2018).

Within the context of ONC, we consider as secondary degeneration all effectors leading to RGC death besides the initial injury (ONC) itself. Following ONC in mice, primary and secondary degeneration cannot be differentiated, as they likely occur simultaneously within the characteristic sigmoidal monophasic loss profile.

Unlike mice, zebrafish exhibit strong neuroprotective properties, showing no RGC loss until 14dpi and only a minor reduction in RGC density by 21dpi. This loss is unlikely to result directly from injury (primary degeneration), as RGC transcriptional profiles at 14dpi closely resemble those of uninjured cells (Zhang et al., 2025). As such, it is more conceivable that this moderate loss of RGCs reflects secondary degeneration. Immune system overactivation is one plausible cause, as immune cell abundance peaks at 7dpi, but only returns to baseline levels by 28dpi (Van houcke et al., 2017). Another possible explanation lies in a recapitulation of developmental mechanisms: during zebrafish retinal maturation, a small proportion of RGCs that fail to form synaptic connections undergo apoptosis (Biehlmaier et al., 2001), a process that also occurs in mice, albeit to a much greater extent (Boia et al., 2020). This mechanism may be reactivated post-injury in adult zebrafish, where regenerating RGCs that fail to reestablish synapses and elicit electrical activity undergo programmed cell death. Therefore, the inability to reform functional connections with target neurons might be another player resulting in secondary degeneration. Notably, this late RGC loss is likely negligible for vision, as zebrafish consistently achieve full functional recovery and re-establish complex visual behaviors after optic nerve injury (Fleisch et al., 2011; Zou et al., 2013; Beckers et al., 2019).

Killifish exhibit strong CNS regenerative capacity in early adulthood, but this ability declines with age, approaching a mammalian-like phenotype at old age (Van houcke et al., 2021; Vanhunsel et al., 2022a, 2022b). Previous efforts to quantify RGC survival post-ONC in killifish relied on biocytin labeling following retrograde tracing using spatially pre-selected cryosections. However, this approach may have introduced variability due to heterogeneous labeling and retinal tissue stretching, respectively. Our whole-tissue analysis builds upon these findings, and aligns with earlier caspase-based studies, which reported peak cell death at 2–7dpi, returning to baseline by 21dpi (Vanhunsel et al., 2022a). Interestingly, both young adult and old killifish display a distinct biphasic RGC loss profile, with a more pronounced second phase of cell loss in older individuals. The mechanisms underlying these distinct kinetics remain unclear but may involve cell-autonomous factors, interactions with the environment of the injured retina, or both. Also, the molecular heterogeneity of RGC types in killifish is still unknown, yet some subtypes may exhibit differential injury susceptibility, akin to mice (Tran et al., 2019), warranting further investigations. Regarding non-cell autonomous mechanisms, both age groups show immune activation post-ONC, with young adults mounting a rapid but transient response that peaks at 2dpi and resolves by 7dpi. In contrast, older individuals exhibit a delayed but more prolonged and intensive response, peaking at 7dpi and resolving by 14dpi (Vanhunsel et al., 2022a). While immune activation may play a role during the first wave of cell death in both age groups, the prolonged and more intense immune response in the old killifish—potentially linked to inflammaging (Vanhunsel et al., 2021; Bergmans et al., 2024)—may help explain the larger magnitude of RGC loss during the second wave. It can, however, not account for the second wave of RGC loss in young adult killifish, where the retina has reached homeostatic levels again by 7 dpi (Vanhunsel et al., 2022a). A possible explanation may involve the impaired reformation of neural circuits: RGCs that do not successfully reconnect and synapse to tectal neurons may not receive the necessary trophic support for long-term survival (Johnson et al., 2009; Claes et al., 2019) or they may eventually undergo programmed apoptosis due to insufficient target reinnervation and synapse formation. Indeed, by 14dpi, young adult and old killifish have only re-established approximately 70% and 30% of their synapses 14dpi, respectively (Vanhunsel et al., 2022a), which may explain the second wave of RGC loss in young adult and the more pronounced decline in old killifish. However, despite the loss of approximately one-third of their RGCs by 21dpi, young adult killifish retain the ability to recover primary vision, an ability aged fish lack (Vanhunsel et al., 2022a). Future research should explore more complex behavioral outcomes, such as social interactions and mating behaviors, to determine the functional impact of the observed RGC loss on vision and fish welfare.

In summary, cross-species comparisons of RGC survival following ONC reveal distinct resilience patterns. Mice exhibit minimal neuroprotection, whereas zebrafish display robust neuroprotective capacities. Killifish, with their biphasic RGC loss profile, potentially offer a unique model to study both intrinsic injury susceptibility (primary degeneration) and cell loss driven by reduced electrical activity and/or detrimental factors in the retinal microenvironment (secondary degeneration). As such, these findings provide a framework for elucidating molecular mechanisms that regulate RGC survival and for identifying potential therapeutic targets to enhance neuroprotection, which could aid in the treatment of optic neuropathies, and by extension of neurodegenerative disorders and traumatic CNS injuries.

## Abbreviations

AMC: age-matched controls
CNS: central nervous system
dpi: days post injury
FWO: fonds wetenschappelijk onderzoek
KDE: kernel density estimation
IQR: interquartile range
Isl2b: ISL LIM Homeobox 2
ONC: optic nerve crush
PBS: phosphate buffered saline
PFA: paraformaldehyde
PID: pre-immune donkey serum
Pou4f1: POU Class 4 Homeobox 1
RBPMS: RNA-binding protein with multiple splicing
Rbpms2: RNA-binding protein with multiple splicing 2
SEM: standard error of mean
WM: whole-mount

## 6. Acknowledgments

The authors would like to thank Simon Buys, Arnold Van Den Eynde, Evelien Herinckx, Véronique Brouwers for the daily animal maintenance and environmental control; Iene Kemps, Lien Andries, Marie Claes and Marijke Christiaens for their technical support. PJS, SB and LuM hold a personal Research Foundation Flanders (FWO, Belgium) fellowship (114525N, 1165020N and 1S42720N, respectively). Experiments are financially supported by KU Leuven (C14/22/074, KAC14/22/074 and KA16-00745) and FWO (G082221N, G092222N).

## 7. Contributions

Conceptualization and experimental design: LM, LuM and SB; Methodology: JDDS, AZ, PJS, LuM and SB; Visualization: LuM and SB; Writing – original draft: JDDS, AZ, SB, LuM; Writing – review and editing: JDDS, AZ, PJS, LM, LuM and SB; Funding acquisitions: PJS, LM, LuM and SB; Project management: LuM and SB; Supervision: LM, LuM and SB

## 8. Supplementary information

**Supplementary table 1:**

Mean RGC count, retinal area and RGC density of non-injured retinas per species and age, along with their relative standard error of the mean (SEM).

**Supplementary table 2**:

Median RGC count, retinal area and RGC density per species, age and timepoint after optic nerve crush (ONC), along with their relative 25^th^ and 75^th^ percentiles. For killifish, the values of age-matched control were included as well.

**Supplementary table 3:**

Results of Kruskal-Wallis two-way ANOVA between injured young adult and old killifish. Multiple comparisons statistics and p-values are reported.

**Supplementary figure 1: Performance metrics of zebrafish and killifish RGC counting models**

**(A)** Linear regression of automated counts versus the average human count for zebrafish retinal ganglion cells (RGCs). The trained model shows very good performance.

**(B)** Bland-Altman plot comparing the counting bias of the two human counters for zebrafish RGCs. The intrinsic difficulty in counting dense RGCs leads to bias across multiple evaluators.

**(C)** Bland-Altman plot comparing the automated count versus the average human count for zebrafish RGCs. The model shows no bias compared to the average human count.

**(D)** Linear regression of automated counts versus the average human count for killifish RGCs. The trained model shows excellent performance.

**(E)** Bland-Altman plot comparing the counting bias of the two human counters for killifish RGCs. The intrinsic difficulty in counting dense RGCs leads to bias across multiple evaluators.

**(F)** Bland-Altman plot comparing the automated count versus the average human count for killifish RGCs. The model shows no bias compared to the average human count.

**Supplementary figure 2: Scaled isodensity map comparison of non-injured mice and fish**

(**A-D**) Scaled isodensity maps of young adult mouse (10-week-old, **A**), young adult zebrafish (21-week-old, **B**), young adult (6-week-old, **C**) and old (18-week-old, **D**) killifish retinal whole-mounts, showing a considerably higher density of retinal ganglion cells (RGCs) in fish species. (D: dorsal, V: ventral, N: nasal, T: temporal)

**Supplementary figure 3: Retinal growth in non-injured zebrafish and killifish**

**(A)** Quantification of retinal area at 21 weeks and at 21 weeks plus 7, 14, and 21days, reveals no significant growth during the experimental time window in zebrafish.

**(B)** Quantification of retinal area in young adult killifish at 6 weeks and at 6 weeks plus 4, 7, 14, and 21days, shows a significant increase in retinal area within the time window of the experiment.

**(C)** Quantification of retinal area in old killifish at 18 weeks and at 18 weeks plus 4, 7, 14, and 21days, does not show significant growth during the experimental time window.

**(D)** Evaluation of retinal ganglion cell (RGC) density in uninjured retinas of killifish along the experimental timeline reveals a significantly lower density of RGCs at 6 weeks plus 21 days compared to 6 weeks, justifying the usage of age-matched control fish (AMC) for quantitative analyses in young-adult animals.

**(E)** Evaluation of RGC density in uninjured (uninj.) retinas of old killifish along the experimental timeline reveals no significant differences in RGC density in the AMC fish.

(**F-H**) Representation of retinal area (**F**), RGC count (**G**)and RGC (**H**) density across all ages and time points examined in killifish. Models were fitted to capture retinal growth over the lifespan, revealing an approximately linear decline in RGC density with age. Data from 1 (**A**) or 2 (**B-H**) independent experiments, presented as median ± 25-75th CI (A-E). Kruskal-Wallis ANOVA. P-values reported within the figure. Polynomial models were fit to the data and the degree. was chosen based on cross-validation. MSE (Mean squared error)

## 9. Data availability

RGCode2 is available on the public GitLab repository

https://gitlab.com/NCDRlab/rgcode2.

