## Supplementary figure 1 for "Differential retinal ganglion cell resilience to optic nerve injury across vertebrate species"

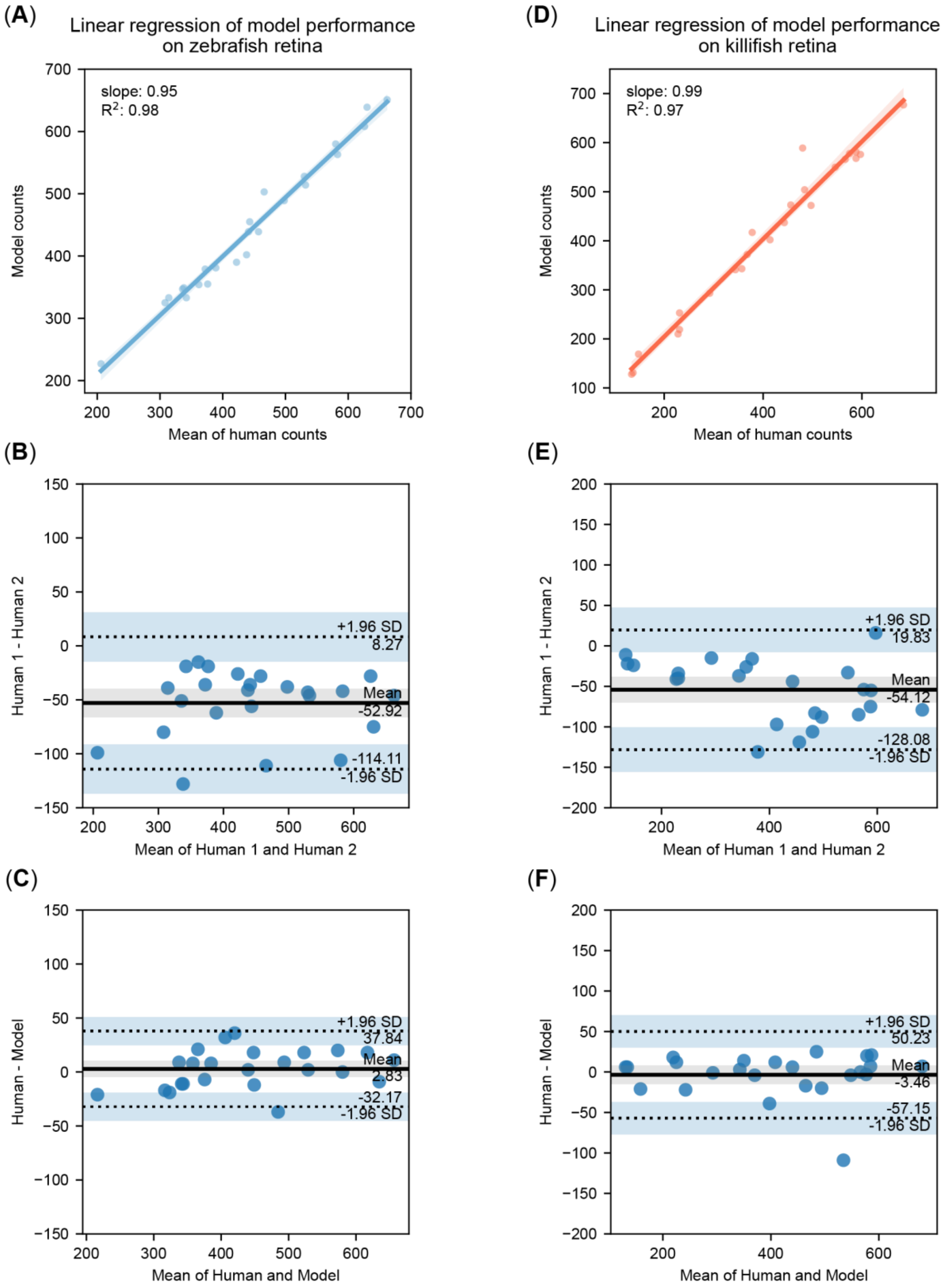


*Supplementary figure 1: Performance metrics of zebrafish and killifish RGC counting models*

(**A**) Linear regression of automated counts versus the average human count for zebrafish retinal ganglion cells (RGCs). The trained model shows very good performance.

(**B**) Bland-Altman plot comparing the counting bias of the two human counters for zebrafish RGCs. The intrinsic difficulty in counting dense RGCs leads to bias across multiple evaluators.

(**C**) Bland-Altman plot comparing the automated count versus the average human count for zebrafish RGCs. The model shows no bias compared to the average human count.

(**D**) Linear regression of automated counts versus the average human count for killifish RGCs. The trained model shows excellent performance.

(**E**) Bland-Altman plot comparing the counting bias of the two human counters for killifish RGCs. The intrinsic difficulty in counting dense RGCs leads to bias across multiple evaluators.

(**F**) Bland-Altman plot comparing the automated count versus the average human count for killifish RGCs. The model shows no bias compared to the average human count.
