## Supplementary figure 2 for "Differential retinal ganglion cell resilience to optic nerve injury across vertebrate species"

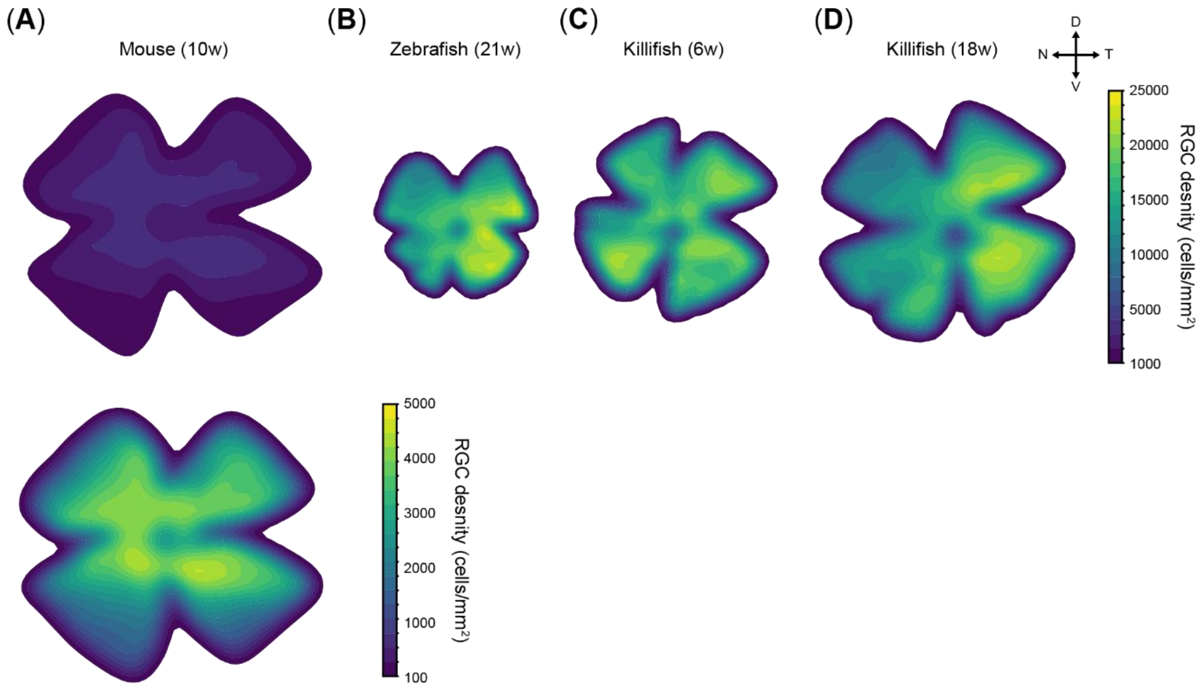


### *Supplementary figure 2: Scaled isodensity map comparison of non-injured mouse and fish*

(**A-D**) Scaled isodensity maps of young adult mouse (10-week-old, **A**), young adult zebrafish (21-week-old, **B**), young adult (6-week-old, **C**) and old (18-week-old, **D**) killifish retinal whole-mounts, showing a considerably higher density of retinal ganglion cells (RGCs) in fish species. (D: dorsal, V: ventral, N: nasal, T: temporal)
