## Supplementary figure 3 for "Differential retinal ganglion cell resilience to optic nerve injury across vertebrate species"

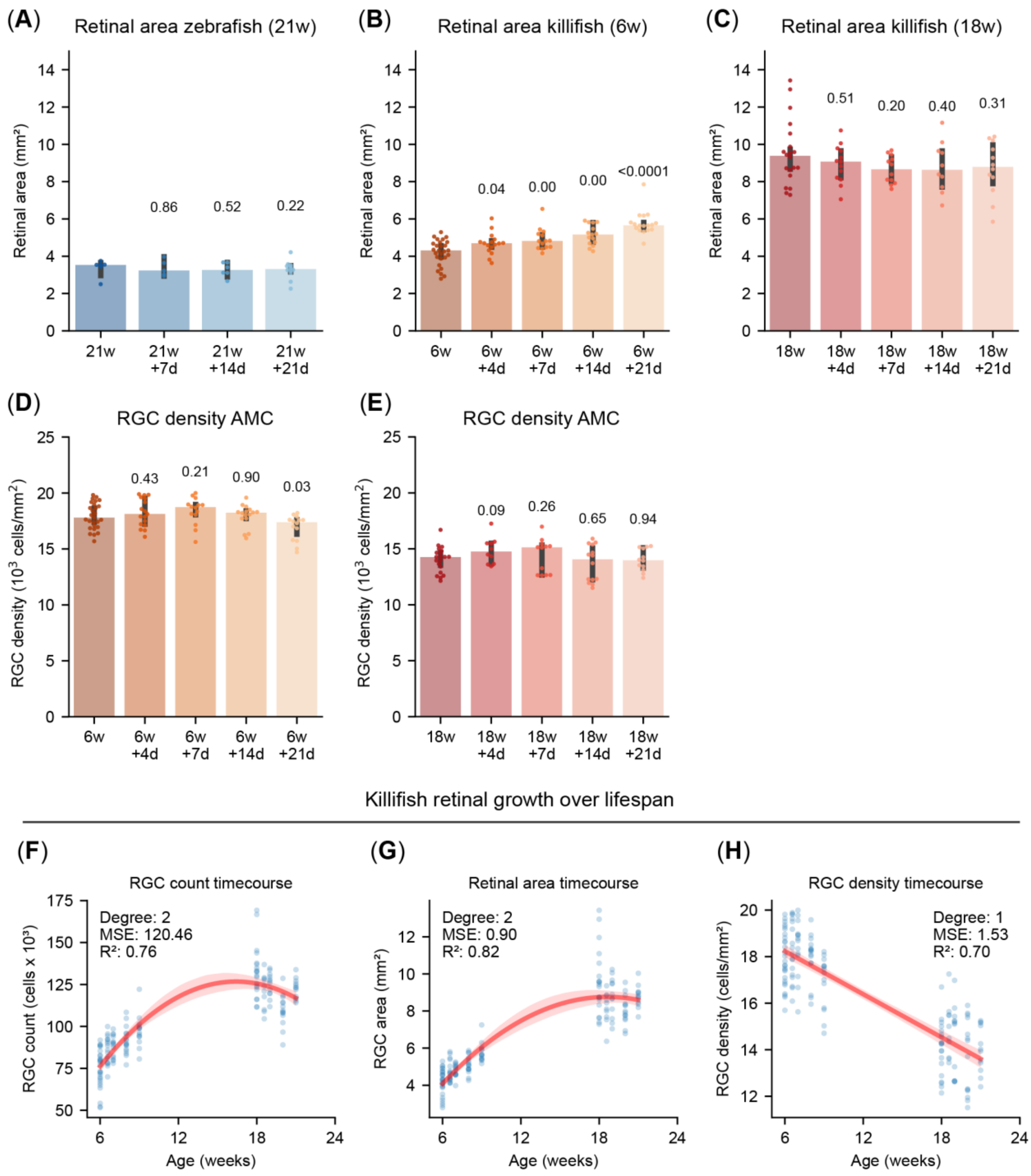


### *Supplementary figure 3: Retinal growth in non-injured zebrafish and killifish*

(**A**) Quantification of retinal area at 21 weeks and at 21 weeks plus 7, 14, and 21days, reveals no significant growth during the experimental time window in zebrafish.

(**B**) Quantification of retinal area in young adult killifish at 6 weeks and at 6 weeks plus 4, 7, 14, and 21days, shows a significant increase in retinal area within the time window of the experiment.

(**C**) Quantification of retinal area in old killifish at 18 weeks and at 18 weeks plus 4, 7, 14, and 21days, does not show significant growth during the experimental time window.

(**D**) Evaluation of retinal ganglion cell (RGC) density in uninjured retinas of killifish along the experimental timeline reveals a significantly lower density of RGCs at 6 weeks plus 21 days compared to 6 weeks, justifying the usage of age-matched control fish (AMC) for quantitative analyses in young-adult animals.

(**E**) Evaluation of RGC density in uninjured (uninj.) retinas of old killifish along the experimental timeline reveals no significant differences in RGC density in the AMC fish.

(**F-H**) Representation of retinal area (**F**), RGC count (**G**)and RGC (**H**) density across all ages and time points examined in killifish. Models were fitted to capture retinal growth over the lifespan, revealing an approximately linear decline in RGC density with age.

Data from 1 (**A**) or 2 (**B-H**) independent experiments, presented as median ± 25-75th CI (A-E). Kruskal-Wallis ANOVA. P-values reported within the figure. Polynomial models were fit to the data and the degree was chosen based on cross-validation. MSE (Mean squared error)
